# Activating alternative transport modes in a multidrug resistance efflux pump to confer chemical susceptibility

**DOI:** 10.1101/2021.12.04.471113

**Authors:** Peyton J. Spreacker, Nathan E. Thomas, Will F. Beeninga, Merissa Brousseau, Colin J. Porter, Kylie M. Hibbs, Katherine A. Henzler-Wildman

## Abstract

Small multidrug resistance (SMR) transporters contribute to antibiotic resistance through proton-coupled efflux of toxic compounds from the bacterial cytoplasm. Previous biophysical studies of the *E. coli* SMR transporter EmrE suggested that it should also be capable of performing proton/toxin symport or uniport, leading to toxin susceptibility rather than resistance *in vivo*. Here we show EmrE does confer susceptibility to several newly characterized small-molecule substrates in *E. coli*, including harmane. *In vitro* experiments show that harmane binding to EmrE triggers uncoupled proton uniport and this protein-mediated dissipation of the transmembrane pH gradient underlies the *in vivo* phenotype. This leads to synergy with some existing antibiotics, such as kanamycin. Furthermore, this shows that it is possible to not just inhibit multidrug efflux but activate alternative transport modes that are detrimental to bacterial growth and metabolism.

## Introduction

There is an urgent need to better understand the underlying mechanisms of antibiotic action and resistance. One mechanism by which bacteria survive antibiotic exposure is through the efflux of toxic compounds by promiscuous multidrug transporters. Most bacterial multidrug resistance transporters operate through an antiport mechanism wherein the efflux of toxic substrates is driven by the downhill import of H^+^ (or Na^+^). As transporters have traditionally been classified as antiporters, uniporters, or symporters, the discovery that the small multidrug resistance (SMR) transporter, EmrE could perform coupled 2 H^+^: 1 substrate (toxin) antiport of a wide range of polyaromatic cations (1) defined its function for many years. However, recent discoveries have highlighted the exceptional promiscuity of transporters in this family despite their small size. The SMR family has become an important model system for studying the mechanism of proton-coupled efflux because of its amenability to structural biology, biophysical experiments, and *in vitro* assays, but this family has repeatedly defied expectations and revealed natural complexity beyond the boundaries of established models. The SMR family was the first membrane protein discovered to have an unusual antiparallel homodimer topology, more recently part of the family was reclassified as toxic metabolite exporters rather than multidrug efflux pumps, and finally, NMR studies of EmrE indicate that this transporter should be capable of multiple modes of transport, not just proton-coupled antiport as required for antibiotic resistance(2–4). Here we focus on the biological implications of alternative transport modes and whether it is possible to not just inhibit multidrug resistance efflux pumps to suppress their contribution to antibiotic resistance, but rather activate alternative transport modes that cause small molecule susceptibility *in vivo*.

Traditional models of proton-coupled antiport focus on the key states and transitions needed for stoichiometric coupled antiport and assume that other states and transitions (leak pathways) contribute minimally to net transport because these alternative pathways would be deleterious in the cell. Recently, careful exploration of the states and transitions of EmrE using NMR revealed that this assumption is incorrect (4). Expanding the mechanistic model to include all the observed states and transitions leads to a more complex free exchange model where proton/toxin symport, proton uniport, and toxin uniport are all theoretically possible in addition to the well-established proton/substrate antiport activity of EmrE (Fig. 1). The biological implications of these alternative transport pathways are significant. While H^+^-driven antiport results in toxin efflux and a resistance phenotype *in vivo*, proton-coupled symport or toxin-uniport would result in active uptake of the toxic molecule into bacteria. Proton leak will rundown the proton motive force, disrupting bacterial energy metabolism, and is also likely to lead to a susceptibility phenotype in bacteria.

**Figure 1:**
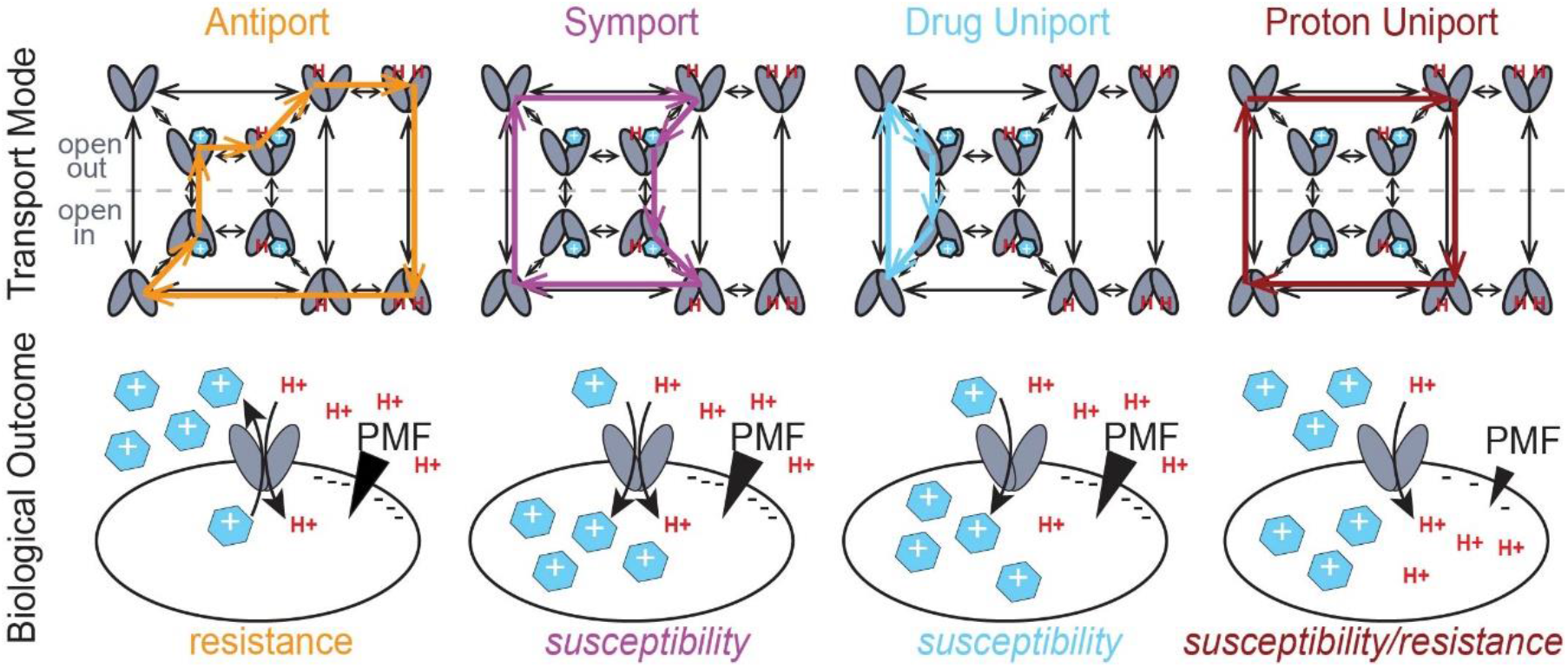
Different transport modes of EmrE result in different biological outcomes. The well-established coupled antiport of proton and drug (orange, top) by EmrE leads to drug resistance *in vivo* (orange, bottom). The Free Exchange Model suggests that EmrE should also be able to perform coupled symport (purple), or drug uniport (blue), either of which would lead to susceptibility rather than resistance *in vivo*. Proton uniport (maroon) could lead to either resistance or susceptibility using an antibiotic adjuvant PMF dissipator with a known antibiotic. The most likely pathway depends on the relative rates of the microscopic steps in the transport cycle, including drug on- and off-rates and the rate of alternating access between open-in and open-out conformations in each state (apo, proton-bound, drug-bound, etc.). Thus, different substrates can lead to different dominant modes of transport and opposing biological outcomes in cells.

There is precedence for converting SMR transporter activity from conferring resistance to susceptibility *in vivo* by mutating the transporter (5). The W63G-EmrE point mutant confers resistance to the clinical antibiotic erythromycin, but susceptibility to polyamine compounds (6), confirming that both transport phenotypes are possible for a single transporter. Of greater biomedical relevance is whether it is possible to shift wild-type (WT) EmrE from its well-established resistance activity to activate alternative transport modes that would confer susceptibility (Fig. 1). There are a few examples of WT transporters utilizing different transport modes to optimize physiological outcomes for sugar uptake under changing external conditions (7, 8) or by preventing loss of acquired metals through back transport (9, 10). EmrE would represent a fundamentally different case where different modes of transport result in *opposite* biological outcomes of resistance versus susceptibility to toxic compounds.

In the case of EmrE, our prior biophysical studies and simulations show that 2 H^+^:1 toxin antiport is kinetically favored under physiological conditions for substrates to which EmrE is known to confer resistance (11). This is in accord with the mechanistic requirement that driving toxin efflux by coupling to the proton motive force, which is inwardly directed in bacteria, requires antiport. However, we have also shown that substrate identity can alter the rate of key microscopic steps in the transport cycle by up to three orders of magnitude (12, 13). Rate changes on this scale have the potential to bias flux through alternative transport pathways and shift the balance of net transport (4). Here we experimentally test whether substrate identity can activate alternative transport modes in WT EmrE. Using an unbiased small molecule phenotypic screen, we identify new substrates to which EmrE confers resistance *and* new substrates to which it confers susceptibility. Harmane is one of the substrates that most strongly activates susceptibility *in vivo*. We use an *in vitro* solid-supported membrane electrophysiology assay to show that harmane triggers uncontrolled proton leak through EmrE, defining the molecular mechanism underlying the susceptibility phenotype. Additional *in vivo* assays confirm that the alternate transport mode activated by harmane acts on the transmembrane pH gradient, and this activity synergizes with kanamycin. This work opens the possibility of activating alternative transport pathways of multidrug transporters to target antibiotic resistance.

## Results

### An unbiased screen reveals new substrates

Previous EmrE substrate screens have focused on quaternary ammonium compounds (QACs) and quaternary cationic compounds (QCCs) commonly transported by multidrug efflux pumps (1, 14–16). To better explore the substrate profile of EmrE, we performed an unbiased screen using the Phenotypic Microarray assay from Biolog, Inc. This screen assesses the impact of diverse compounds on *E. coli* metabolic output in a differential comparison of MG1655Δ*emrE E. coli* expressing either wildtype or non-functional EmrE (E14Q-EmrE). If the metabolic output was greater when wildtype EmrE was expressed, it indicates that functional EmrE is beneficial, and the compound was classified as a resistance hit. If the metabolic output was greater when E14Q-EmrE was expressed, it indicates that functional EmrE is detrimental and the compound was classified as a susceptibility hit (Fig. 2, Fig. S2, Table S1, see methods for selection criteria). As shown in Fig. 2A, the screen identified compounds in both categories: resistance and susceptibility.

**Figure 2:**
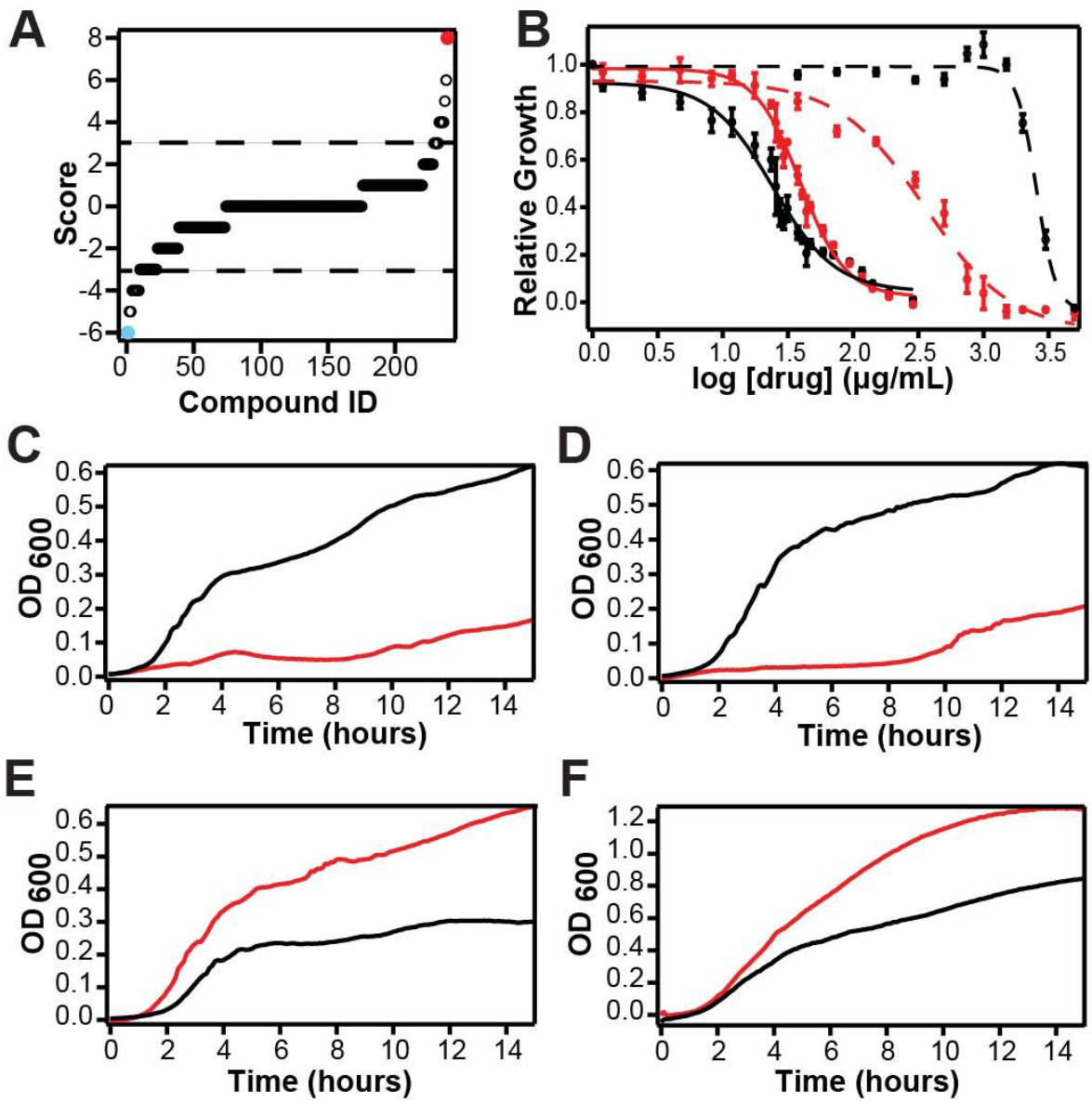
EmrE can confer either resistance or susceptibility *in vivo*. (A) Biolog phenotype microarray results were sorted by hit score for all 240 compounds in the screen (see Methods). Scores above +3 or below -3 are considered resistance or susceptibility hits, respectively, based on the differential between functional (WT) and non-functional (E14Q-EmrE). The strongest resistance hits (red) and susceptibility hits (cyan) were tested in growth assays (C-F). (B) IC_50_ curves of WT- (black) and E14Q-EmrE (red) are shown for ethidium bromide (dashed lines, resistance) and harmane (solid lines, susceptibility). Note that cells expressing WT-EmrE have a 40% lower IC_50_ value than cells expressing E14Q-EmrE in the presence of harmane. MG1655 ΔemrE *E. coli* expressing WT-EmrE (black) or E14Q-EmrE (red) were grown in the presence of (C) 0.5 mM methyl viologen (MV^2+^), (D) 0.05 mM chelerythrine chloride (CC), (E) 0.1 mM 18-crown-6-ether, or (F) 0.13 mM harmane. As expected, *E. coli* expressing WT-EmrE survived in the presence of MV^2+^ and CC (C, D), but *E. coli* expressing E14Q-EmrE did not, consistent with a resistance phenotype. In contrast, *E. coli* expressing non-functional, E14Q-EmrE had a higher OD_600_ at the stationary phase than *E. coli* expressing WT-EmrE in the presence of 18-crown-6-ether and harmane, (E, F), consistent with a susceptibility phenotype.

The well-established EmrE substrate methyl viologen (MV^2+^) was the strongest resistance hit with the highest possible score according to our criteria (+8). Acriflavine, another known substrate, was also a strong +4 resistance hit, confirming that the Biolog assay accurately reports on EmrE toxin resistance phenotypes. Chelerythrine chloride has not been previously identified as an EmrE substrate but showed a strong +5 resistance phenotype. Microplate growth assays of *E. coli* expressing either wildtype or E14Q-EmrE in the presence of MV^2+^ or chelerythrine chloride confirmed the resistance phenotype (Fig. 2C, D, Fig. S3). Chelerythrine chloride has been used as an antibacterial agent for toxin-resistant infections (17), so EmrE-conferred resistance may be clinically relevant.

EmrE’s resistance activity has been well characterized in *E. coli*, but a susceptibility phenotype for WT EmrE has not previously been reported. The top three susceptibility hits identified in the Biolog screen were: harmane (−6), hexachlorophene (−6), and 18-crown-6-ether (−5). Compared to the other susceptibility hits, hexachlorophene is extremely insoluble therefore it was not evaluated further. In microplate growth assays, E14Q-EmrE expressing cells had a higher final OD_600_ at stationary phase in the presence of 18-crown-6-ether (Fig. 2E, red line) or harmane (Fig. 2F, red line), but cells expressing WT EmrE had significant growth deficiencies after five hours of treatment (black lines), manifested by an earlier onset of stationary phase with lower OD_600_. This confirmed that functional EmrE confers susceptibility rather than resistance to these compounds. 18-crown-6 ether has previously been implicated in cellular toxicity due to interference with cation transport (18–20), but the mechanism of the possible antimicrobial activity of harmane is unknown (21, 22).

### Harmane binds EmrE and triggers uncoupled proton transport

We acquired ^1^H-^15^N HSQC NMR spectra of WT-EmrE in the presence and absence of harmane (Fig. 3A) to determine whether a direct binding interaction between EmrE and harmane could be responsible for the susceptibility phenotype. The NMR chemical shift perturbations (CSPs) observed for a subset of peaks upon harmane binding indicate that there is direct interaction between EmrE and harmane at a localized binding site. In the absence of substrate, alternating access of EmrE occurs on the intermediate timescale leading to many broadened peaks (black spectrum). Upon harmane binding, additional peaks are resolved in locations where resolved peaks are observed for EmrE bound to other substrates with experimentally established alternating-access rates in the slow-exchange regime (23), indicating that harmane binding likely slows the rate of alternating access in EmrE. The extensive dynamics in the drug-free state preclude backbone assignment and residue-specific CSP calculation. Direct binding was further confirmed using intrinsic tryptophan fluorescence quenching (12) (Fig. 3B).

**Figure 3:**
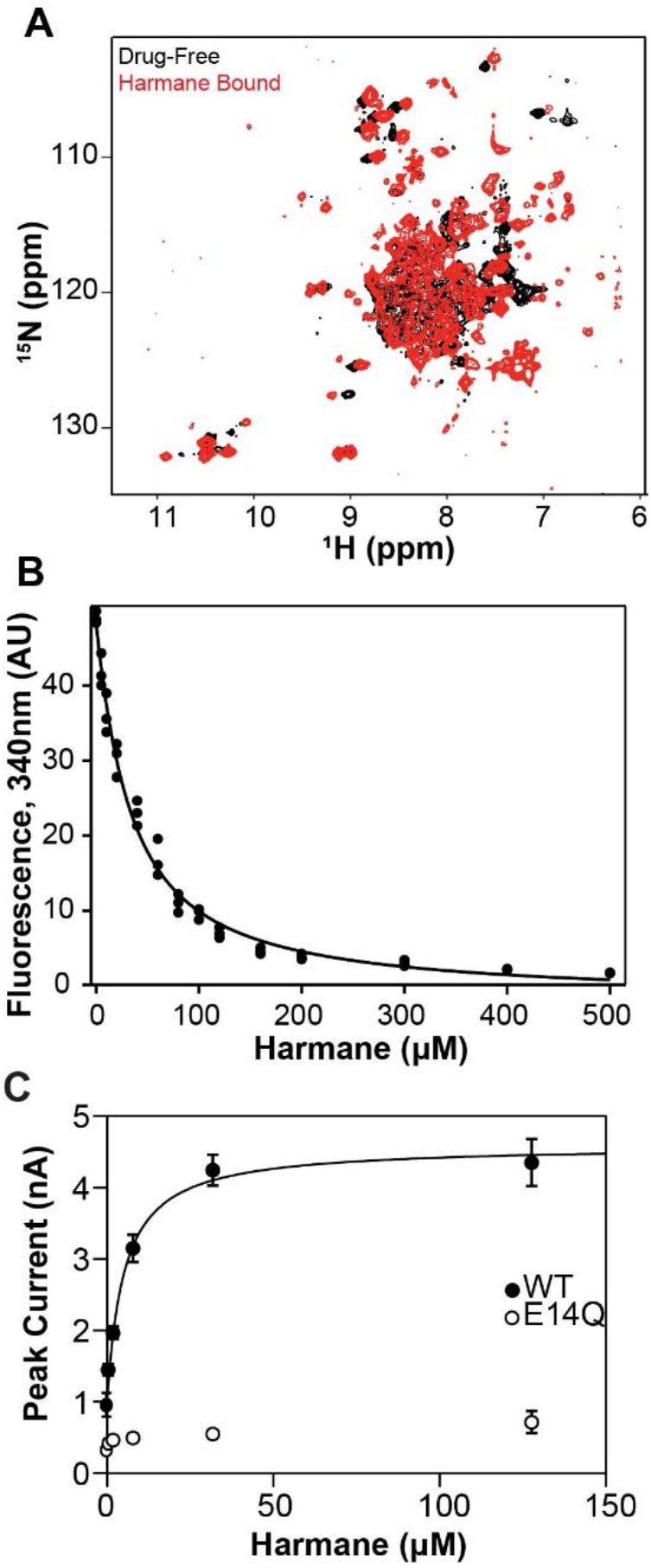
EmrE directly binds harmane *in vitro*. A) ^1^H^15^N-TROSY-HSQC spectra of WT EmrE in isotropic bicelles (q=0.33) at 45°C without (black) or with (red) harmane bound. B) Harmane quenches intrinsic tryptophan fluorescence of WT-EmrE in a dose-dependent manner, with an apparent K_d_ value of 29 ± 2μM. Together, A and B confirm that harmane binds WT-EmrE directly at a specific site. C) SSME measurement of EmrE transport current shows that peak current increases with harmane concentration and saturates, consistent with transport dependent on harmane binding. The same 2-fold proton gradient (pH 7.0 outside, pH 7.3 inside) was used as in Figure 4. The transport fits to Michaelis-Menten kinetics with a K_m_ value of 5 ± 1μM. The non-functional control (E14Q) displayed no significant current at any concentration of harmane, confirming that harmane-induced proton-flux through EmrE requires functional transporter.

To determine whether harmane binding triggers EmrE transport activity, we turned to solid supported membrane electrophysiology (SSME). SSME allows the detection of net charge movement in proteoliposomes adsorbed onto a gold electrode sensor upon buffer perfusion and is ideal for measuring small transport currents produced by moderate-flux transporters such as EmrE (24). Harmane triggered EmrE transport currents, which increased and eventually saturated with increasing harmane concentrations (Fig. 3C). While this data is strongly suggestive of a direct effect of harmane on the transport activity of EmrE, how might EmrE transport lead to drug susceptibility?

EmrE-mediated drug resistance phenotypes can only be explained by the canonical proton/drug antiport mechanism, but toxin susceptibility can arise from three potential transport mechanisms: drug uniport, proton uniport, or proton/drug symport (Fig. 1). To better understand how EmrE confers susceptibility to harmane, we performed additional SSME experiments using an assay recently developed in our lab to characterize the ion-coupling behavior of secondary active transporters (25, 26).

The hallmark of coupled transport is the ability of downhill transport of one substrate to drive uphill transport of another substrate. The difference between antiport and symport is simply which orientation of the drug gradient (relative to the smaller proton gradient) enhances proton-driven transport and which orientation reverses net transport. In the SSME assay, transport is initiated by buffer perfusion to create substrate gradients across the liposomal membranes. Various combinations of substrate gradients (Fig. 4A) will have different and predictable effects on the transport signal in the case of antiport, symport, drug uniport, or proton uniport (Fig. 4B).

**Figure 4:**
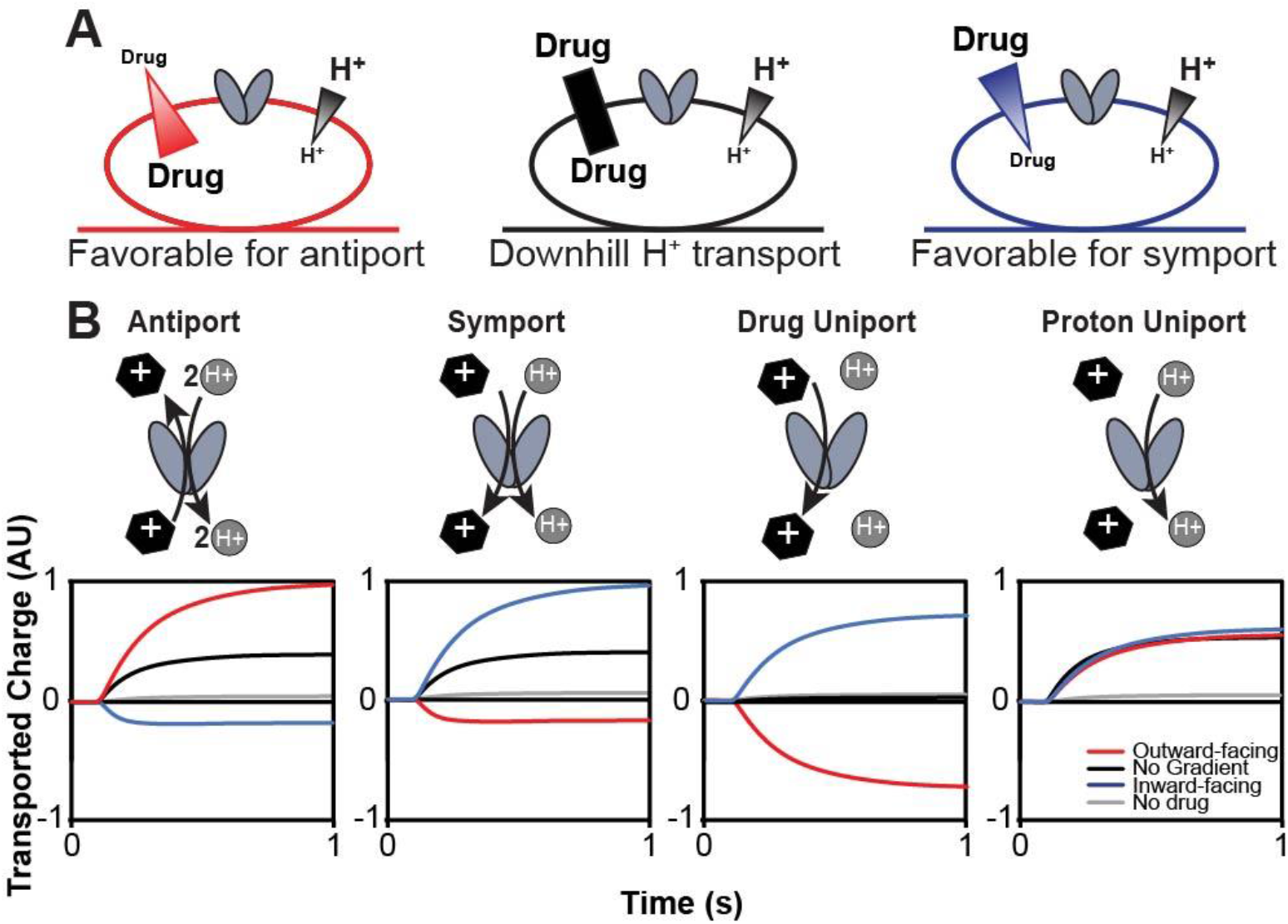
Determination of transport mode *in vitro* using solid-supported membrane electrophysiology (SSME). Expected results in the SSME assay for the different transport modes of EmrE as predicted by the Free Exchange Model. A) Three conditions are used to determine the transport mode of EmrE using SSME. In all cases, there is a two-fold inward-driven proton gradient (pH 7.0 outside, pH 7.3 inside). To test for antiport (red), a 16-fold substrate gradient was pointed against the proton gradient. For symport (blue), the 16-fold drug gradient is oriented with the proton gradient. A third condition testing the presence of a drug without a gradient (black) acts as a control for drug-activated proton uniport.

In the absence of a drug gradient (black), transport is driven by the two-fold inward-facing proton gradient, resulting in a positive signal for the canonical 2 H^+^/1 drug^+^ antiport (net +1 inward per transport cycle), symport (net +1 or +2 inward, for H^+^ and a neutral or drug^+^), or proton uniport (net +1 inward). For drug uniport, a proton gradient alone will not drive transport, resulting in no signal. In the case of drug/proton antiport, the addition of a much larger drug gradient opposite the proton gradient (red) will favor antiport and cause a larger positive signal, while aligning the drug-and proton-gradients in the same direction (blue) requires one substrate to move against its concentration gradient. Under our experimental conditions, the driving force from the drug gradient out-competes the proton gradient and reverses the direction of net transport compared to the 2-fold proton gradient alone. In contrast, uncoupled transport depends solely on the gradient of the uniported substrate. Drug uniport depends only on the direction of the imposed drug gradient and the net charge on the drug (shown for drug^+^, no current would be observed under any condition for an uncharged substrate such as harmane). Proton uniport will result in a consistent, positive signal due to proton flux down the uniform two-fold proton gradient under all three conditions. Transport should be minimal in the absence of drug (gray) as EmrE does not spontaneously leak protons (4).

We first measured net charge movement under the three gradient conditions depicted in Fig. 3A for the transport of methyl tetraphenylphosphonium (MeTPP^+^), which is known to be antiported by EmrE (Fig. 5A, B; Fig. S4A, B). Proteoliposomes reconstituted with E14Q-EmrE (dashed lines) were used as negative controls and produced minimal signals under all conditions, regardless of substrate. In the absence of drug, the proton gradient alone induces a small positive current in WT EmrE proteoliposomes, indicating minimal proton leak. When MeTPP^+^ is added, we observe transport reversal as expected for 2 H^+^ / 1 MeTPP^+^ antiport.

**Figure 5:**
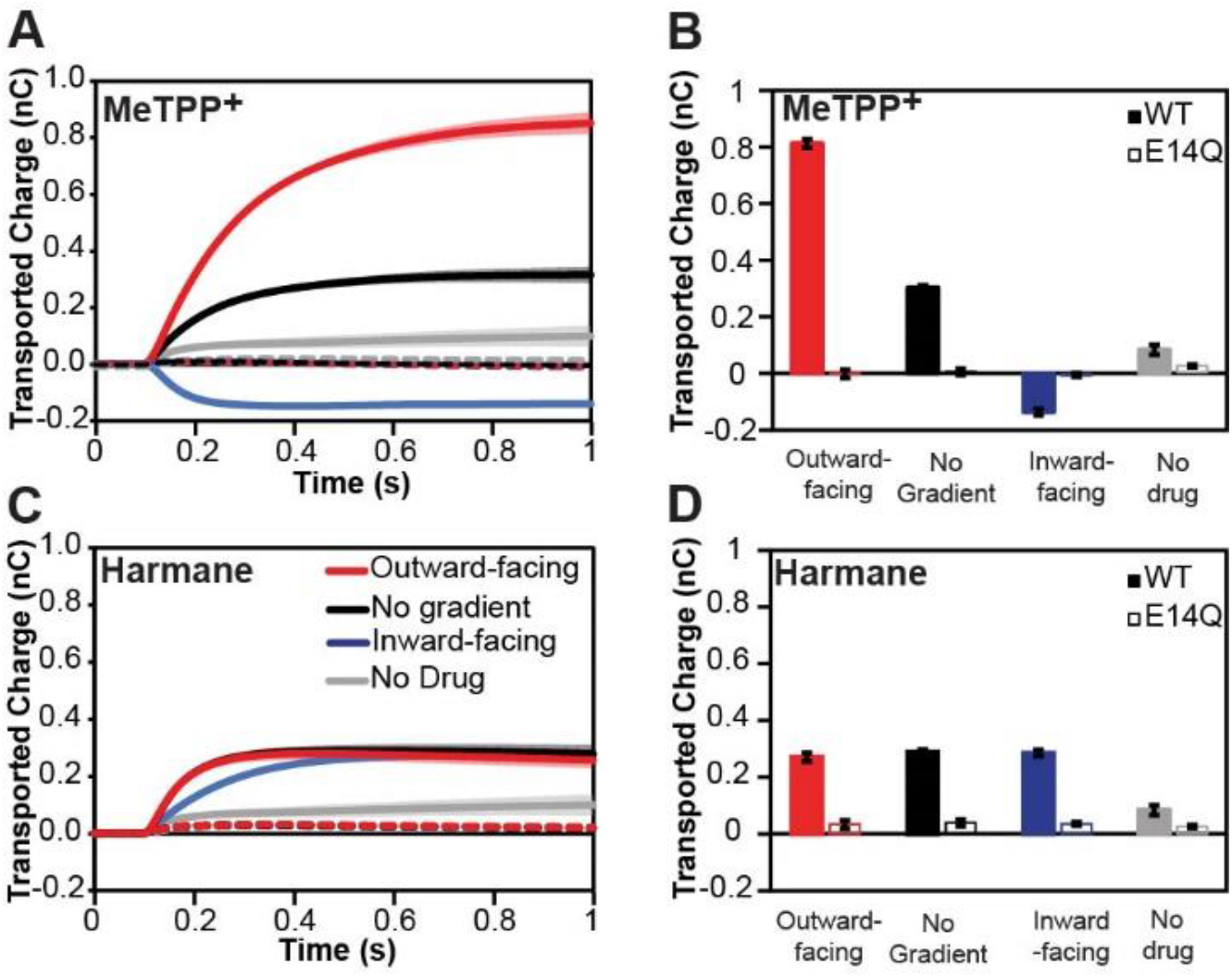
Harmane induces proton flux through EmrE. MeTPP^+^ (A, B) behaves as expected for an antiported substrate (see Figure 3). The total transported charge reverses when the drug gradient is inverted, characteristic of coupled antiport. In contrast, the harmane transport signal (C, D) is the same regardless of the harmane gradient, matching the expected behavior for downhill proton transport (proton leak). The current is minimal in the absence of drug (B, C; gray) or for liposomes containing non-functional E14Q-EmrE (dashed lines), indicating that the observed charge transport is due to substrate-triggered EmrE activity.

In contrast, net transport does not reverse when harmane is the substrate (Fig. 5C, D; Fig. S4C, D). Instead, the net charge transport is constant and positive (down the proton gradient), under all conditions. This lack of reversal between inwardly and outwardly directed drug gradients is indicative of uncoupled proton transport (leak). This proton leak is triggered by harmane since the signal is larger with harmane than the background proton leak in the absence of drug.

### Harmane dissipates the ΔpH component of the proton motive force

Reexamining the cell growth assays indicates the significant harmane phenotype appears around the 5-hour mark (Fig. 2E, F), approximately the point at which fermentable sugars are consumed (27) and cells become more reliant on the proton motive force for ATP production when *E*.*coli* are grown in LB. Thus, the *in vitro* and *in vivo* assays so far are consistent with harmane triggering an EmrE-mediated uncontrolled proton leak. To explore the mechanism of proton leak in bacteria, we performed checkerboard assays with kanamycin and tetracycline and measured growth curves and IC_50_ values with co-treatment of harmane and bicarbonate.

Bicarbonate acts to dissipate the ΔpH component of the proton motive force (28). Consequently, we expect that the harmane phenotype would be reduced in the presence of bicarbonate since both drugs act on the ΔpH. We therefore determined the IC_50_ of MG1655 Δ*emrE* cells expressing WT-or non-functional EmrE treated with harmane and bicarbonate simultaneously. The IC_50_ for harmane in the presence of 25 mM sodium bicarbonate (Fig. 6C) cannot be determined quantitatively because it is beyond the solubility limit of harmane in solution. In contrast, the harmane IC_50_ determined in the absence of bicarbonate (Fig. 2B), is 1.1 mM. The antagonism between bicarbonate and harmane is consistent with harmane dissipating the ΔpH component of the proton motive. In addition, the differential effect of harmane on cells expressing WT versus non-functional EmrE is clear in Figure 2B, reflecting that harmane acts via EmrE. However, when bicarbonate is added there is no longer a significant difference with functional versus non-functional EmrE, consistent with ΔpH dissipation by bicarbonate in an EmrE-independent manner. These results support the *in vitro* assays indicating harmane triggers uncoupled proton leak through EmrE and dissipates the ΔpH component of the proton motive force through EmrE *in vivo*.

**Figure 6:**
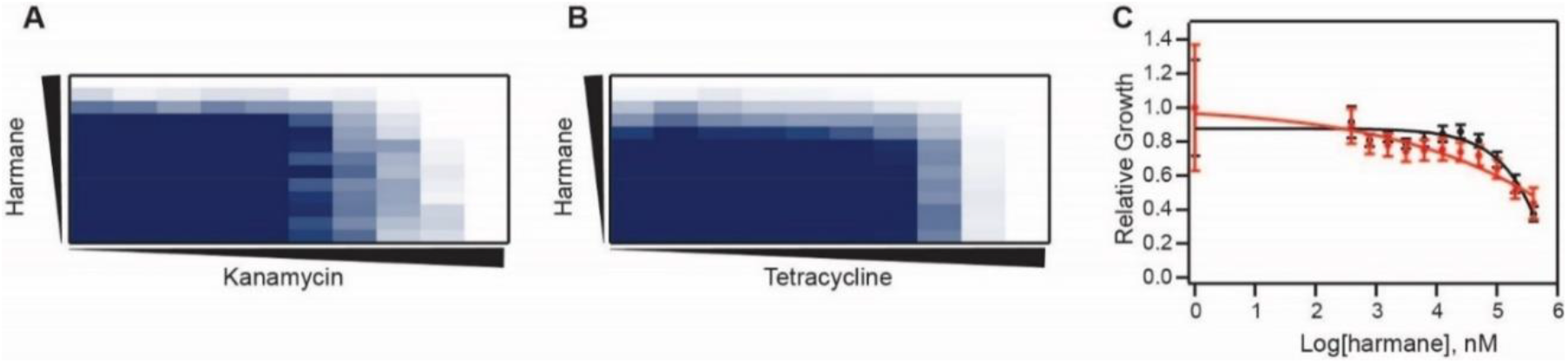
Harmane dissipates the proton gradient bacterial cells. MG1655 Δ*emrE E. coli* expressing WT-EmrE were grown in the presence of harmane as an adjuvant with kanamycin (A) and tetracycline (B). The results shown are the average of three biological replicates. In the presence of harmane, it takes less kanamycin to reach the 10% growth mark as indicated by the step pattern in A. This pattern is not seen when harmane is added as an adjuvant to tetracycline. Further, the mean FIC values of the experiments are 0.375 (synergistic), and 0.58 (indifferent) for kanamycin and tetracycline, respectively. Compound concentrations are denoted by the black triangles increasing left to right for kanamycin and tetracycline and bottom to top for harmane. (C) Relative growth was plotted as a function of harmane concentration in the presence of 25 mM sodium bicarbonate for MG1655 Δ*emrE E. coli* cells expressing WT-(black) and nonfunctional, E14Q-EmrE (red). Compared to the results from Figure 2B, bicarbonate diminishes the susceptibility phenotype in *E. coli* expressing WT-transporter. Growth is shown as a gradient of highest OD_600_ (navy) to no growth (white).

Checkerboard assays are useful for assessing the synergy or antagonism of drugs that act on the proton motive force and determining which of the two components, the proton gradient (ΔpH) or the charge gradient (ΔΨ), the drug primarily targets (29, 30). The aminoglycoside kanamycin requires the ΔΨ component of the proton motive force to cross the inner membrane and reach its cytoplasmic targets. Conversely, tetracycline requires the ΔpH and would produce an antagonistic interaction if the drug being tested dissipates the ΔpH component. If the drug of interest dissipates the ΔpH, as demonstrated for harmane *in vitro*, the ΔΨ component of the proton motive force will increase to compensate as part of normal bacterial homeostasis. This will lead to a stronger ΔΨ and therefore more efficient uptake of kanamycin – leading to synergy and a reduction in the kanamycin minimum inhibitory concentration (MIC). The mean fractional inhibitory concentration (FIC) index was calculated and used to determine synergism (FIC < 0.5), indifference (0.5 ≤ FIC < 1), or antagonism (FIC ≥ 1).

Figure 6 shows the results of checkerboard assays with harmane and kanamycin or tetracycline. These assays were performed in MG1655 Δ*emrE E. coli* expressing WT-or non-functional EmrE to confirm that harmane-triggered EmrE activity *in vivo* is due to ΔpH dissipation. In the harmane/kanamycin experiment (Fig. 6A), a characteristic stair pattern is observed as expected for a synergistic relationship between the two compounds. An FIC value of 0.37 was determined similar to previous reports (29) for kanamycin and harmane, suggesting synergy. When the same analysis was performed with tetracycline (Fig. 6B), an FIC value of 0.58 was determined, indicating indifference between tetracycline and harmane. No synergy is observed when *E. coli* expressing non-functional transporter are used (Fig. S5). In addition, these results demonstrate that small molecules like harmane that trigger proton leak through EmrE may synergize with certain classes of antibiotics with independent modes of action.

## Discussion

SMR transporters are found throughout the bacterial kingdom and efflux toxic hydrophobic cations through coupled antiport of substrate and protons as illustrated in Figure 1 (31–34). The most widely studied member of this family, EmrE, confers resistance to a broad array of toxic polyaromatic cations in *E. coli* (16, 35). Although this small protein was originally proposed to be an ideal model for studying the molecular mechanism of proton-coupled antiport, detailed biophysical studies have instead revealed surprising promiscuity not just in substrate specificity, but also in transport mechanism. The data presented here confirm that alternative transport modes suggested by that prior work can be activated *in vivo* by small molecules, such that some small molecules trigger EmrE-mediated susceptibility rather than resistance. In particular, the susceptibility-activating substrates we have discovered thus far trigger uncontrolled proton leak through EmrE and dissipate the transmembrane pH component of the proton motive force.

Targeting bacterial bioenergetics as an alternative to cell envelope biogenesis or macromolecular biosynthesis is an area of active interest for novel antibiotic development (29, 30, 36, 37) as well as synergy or collateral susceptibility with current antibiotics (38– 41). While several PMF-targeting molecules are available such as gramicidin or nigericin, they generally act non-specifically by creating pores in the lipid bilayer. In other cases, such as the recently identified antimicrobial halicin, the PMF is targeted through an unknown mechanism (30). The results presented in this paper demonstrate that small molecules can run down the proton motive force by triggering alternative transport modes in a small multidrug resistance transporter to a level that is detrimental to bacterial growth and metabolism. The *in vivo* data demonstrate that this phenotype is detrimental to cell growth and metabolism and can synergize with the activity of existing antibiotics that utilize other components of the proton motive force.

Kinetic studies of purified transporters show that some transporters appear to be tightly coupled and highly efficient while others are more loosely coupled, although the experimental challenges of performing these experiments have limited the number of transporters whose mechanism has been rigorously characterized. More recently, there has been renewed interest in kinetic modeling to understand how these more complex network models still achieve relatively efficient coupled transport and may be important for optimizing overall biological function (11, 42, 43). In ATP-coupled transport systems, a more significant “leak” (uncoupled ATP hydrolysis) is observed for promiscuous transporters than for highly selective transporters. For example, the multidrug efflux pump P-glycoprotein exhibits significant levels of basal ATP hydrolysis (44). Loose coupling between the driving force, whether that consists of an electrochemical ion gradient or ATP hydrolysis, and substrate transport may be a requirement of multidrug recognition and efflux, as tight binding generally requires highly specific and selective interactions between the protein and the substrate. The possibility that loose coupling would extend to ion-coupled multidrug transporters, including the SMR family, was originally discussed more than 20 years ago (45). Here we show that a small molecule can exploit this property of promiscuous multidrug transporters and trigger protein-mediated proton leak. If loose coupling is required for multidrug efflux, targeting dissipative pathways in multidrug transporters may represent a new general strategy for combatting antibiotic resistance, either through the development of novel proton-motive-force-dissipating antibiotics or in combination to restore the efficacy of current antibiotics.

## Materials and Methods

### Plasmids and Strains

All *in vivo* experiments were performed in MG1655 Δ*emrE E. coli* cells (Item number: JW0531-2, *E. coli* Genetic Resource (CGSC), Yale) transformed with a low copy number plasmid under a pTrc promoter. *In vivo* experiments relied on leaky expression of these plasmids and expression levels were validated by western blot analysis (Fig. S1). Protein expression utilized BL21 (Gold) DE3 *E. coli* transformed with a pET15b plasmid containing the respective EmrE construct. Detailed validation of the pWB plasmid expression as well as vector controls can be found in Fig. S1. Expression levels were validated from the pWB plasmid by western blot using an anti-His HRP conjugate kit (Qiagen) (Fig. S1).

### Biolog Phenotypic Microarrays

MG1655 *ΔemrE E. coli* cells expressing either WT-or E14Q-EmrE constructs were grown on lysogeny broth media with ampicillin overnight at 37°C. The phenotype microarray tests followed the established protocols of standard phenotype microarray (PM) procedures for *E. coli* and other gram-negative bacteria (12). PM01-20 plates were used to screen both WT-and E14Q-EmrE expressing *E. coli* ([Biolog website](https://www.biolog.com/products-static/phenotype_microbial_cells_overview.php)). Overnight plates were resuspended in IF-0a inoculating fluid (Biolog) to an optical density of 0.37. The cells were diluted by a factor of 6 into IF-0a media plus Redox Dye A and 20mM glucose was added for PM3-8 plates. Cells were diluted to a 1:200 dilution in IF-10a media (Biolog) with Redox Dye A for PM9-20 plates. PM plates were inoculated with 100µL of cell suspensions per well. The microplates were incubated at 37°C and read using the OmniLog instrument every 15min for 24 h. The area under the resulting metabolic curves was determined for cells expressing WT-EmrE or E14Q-EmrE. The difference was calculated using the equation:

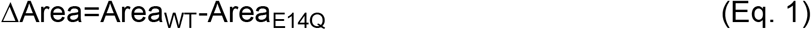

This equation resulted in positive values for greater respiration by cells expressing WT-EmrE and negative values for greater respiration by cells expressing non-functional EmrE. The 10% trimmed mean was then calculated for each data set (WT replicate 1, WT replicate 2, E14Q replicate 1, E14Q replicate 2) separately for each transporter as variation between replicates can arise due to minor deviations between plate sets or in the exact concentration of dye or OD of cells upon dilution on different days. The standard deviation was then calculated among known non-hits (selecting at least 50 wells out of the 960 total wells in a single data set) to determine the cut-off values for actual hits. Individual wells were assessed as hits if the calculated Delta value (equation 1) was more than two standard deviations from the 10% trimmed mean. For each hit, a value of +1 was assigned for resistance hits (positive Delta), and a value of -1 was assigned for susceptibility hits (negative Delta). These values were then summed across all eight wells for a single compound (4 wells of the same compound per plate set * 2 replicates, with a max score of ±8. Final resistance or susceptibility hits were assigned if the total score was ≥ +3 (resistance) or ≤ -3 (susceptibility). This definition was chosen since small total hit scores of ±1 or ±2 could arise by chance using the ± 2*SD cutoff to score individual wells. Values of ±3 recognize consistent hits across multiple replicates and/or different concentrations of the same compound. Our cutoff is not set higher since the 4 wells of each compound on a single plate set include different concentrations and some concentrations may not be sufficient to elicit a phenotype.

### Microplate Growth Assays

Cells expressing plasmids of interest were grown in Mueller-Hinton broth (Sigma, 100µg/mL ampicillin, pH 7.0) from a single colony to an OD of 0.2 at 37 °C. The cells were then diluted to a final OD of 0.01 in 384-well microplates containing concentration ranges of MV^2+^, harmane, 18-crown-6-ether, and chelerythrine chloride. The plates were incubated and shaken in a microplate reader (BMG-Labtech) at 37°C. OD_600_ was measured every 5 minutes for 20 hours. Experiments were performed with four biological replicates and data were analyzed using Igor Pro v8 (WaveMetrics Inc.).

### IC_50_ Value Determination

MG1655 Δ*emrE E. coli* cells expressing either WT or E14Q-EmrE were grown overnight at 37 °C from a single colony. Concentration ranges of ethidium bromide (0-5 mM) and harmane (0-0.4mM) were assayed in microplates with a starting OD_600_ of 0.1. Plates were then incubated with shaking for 18 hours with shaking at 37 °C. OD_600_ endpoints were taken using a BMG plate reader. Relative growth was calculated by dividing the measured OD_600_ from a given concentration by the OD_600_ for cells containing no s. Experiments were performed in triplicate and fit a simple sigmoid equation using Igor Pro v8 (WaveMetrics Inc.).

### EmrE expression and purification

BL21 Gold (DE3) *E. coli* cells were transformed with pET15b-EmrE, pET15b-S64VEmrE, or pET15b-E14QEmrE plasmids and grown in M9 minimal media to an OD_600_ of 0.9. The bacteria were flash cooled and then induced with 0.33M IPTG overnight at 17 °C. The *E. coli* cells were collected with centrifugation, lysed, the membrane fraction solubilized with decylmaltoside (DM), and the proteins purified using nickel affinity chromatography followed by size exclusion chromatography (SEC) on a Superdex 200 column as previously described(13). Protein concentrations were determined using absorbance at 280 nm with an extinction coefficient of 38,400 L/mol cm(46).

### Intrinsic Tryptophan Assays

Purified WT- and S64V-EmrE were reconstituted into isotropic bicelles of DMPC/DPHC (q=0.33) as previously described(47). Reconstitution of purified EmrE into liposomes was performed as described above for SSME transport assays but using DMPC lipids with an EmrE:DMPC ratio of 1:75. Bicelle stocks (2X) were prepared by dissolving DMPC in assay buffer containing 100 mM MOPS pH 7.0, 20 mM NaCl to a final concentration of 300 mM and incubating at 45 °C for 1.5 hrs. DHPC was then added to a final concentration of 100 mM to create q=0.33 isotropic bicelles, incubated an additional hour, and subjected to 3 freeze/thaw cycles. Harmane was prepared to a maximal concentration of 800 μM in assay buffer with 1X bicelle stock and rotated for 72 hours then serial diluted into black 96-well flat-bottom plates. WT- and S64V-EmrE in DMPC/DHPC bicelles were added to a final dimer concentration of 10 μM and the plate was incubated at room temperature for one hour. The final assay volume was 200 μL and each concentration was present in triplicate. Endpoint fluorescence was determined using a TECAN Spark and data analysis was performed in Igor Pro v8. Data were fit to a single binding isotherm detailed in the following equation:

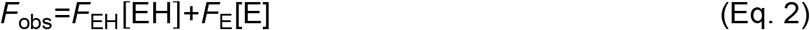

Where *F*_obs_ is the observed fluorescence, *F*_EH_ is the fluorescence of the EmrE functional dimer bound to harmane, [EH] is the concentration of EmrE functional dimer bound to harmane, *F*_E_ is the fluorescence of the EmrE functional dimer, and [E] is the concentration of EmrE functional dimer.

[EH] is calculated from the following equation:

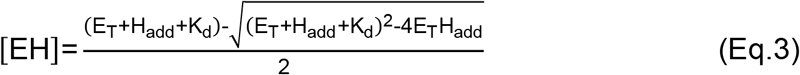

Where E_T_ is the total concentration of EmrE functional dimer in the sample, H_add_ is the total added harmane in the sample, and *K*_d_ is the dissociation constant.

The concentration of unbound EmrE functional dimer ([E]) is given by the following equation:

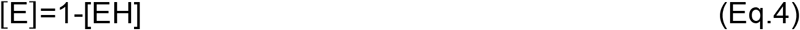

### Direct binding by NMR spectroscopy

Purified ^15^N^2^H-EmrE (0.7-1 mM) was reconstituted into isotropic bicelles (q=0.33) at pH 4.5. The harmane-bound EmrE sample was soaked in harmane overnight with incubation at 45°C. HN-Transverse relaxation optimized spectroscopy – heteronuclear single quantum correlation (HN-TROSY-HSQC) experiments were performed on a 750MHz Bruker Avance spectrometer at 45°C (d1 = 2 sec). Spectra were processed and analyzed using NMRPipe and CCPnmr Analysis 3.0.4.

### Solid Supported Membrane Electrophysiology Transport Assays

WT- and E14Q-EmrE were expressed and purified, with the final SEC performed in assay buffer (50 mM MES, 50 mM MOPS, 50 mM bicine, 100 mM NaCl, 2 mM MgCl_2_, 40 mM DM, pH 7). All buffers were carefully adjusted to the desired pH exclusively with NaOH to ensure consistent Cl^-^ concentrations across the membrane for transport assays. Protein was reconstituted into 1-palmitoyl-2-oleoyl-sn-glycero-3-phosphocholine (POPC) proteoliposomes at a lipid-to-protein ratio of 1:400 in pH 7 assay buffer. Briefly, 15 mg/ml stocks of POPC were diluted in assay buffer and incubated at 45 °C for 1 hour. Lipids were bath sonicated for 1 min then octyl glucoside (OG) was added to a final concentration of 0.5%. Lipids were sonicated for an additional 30 seconds and returned to 45 °C to incubate for 15 minutes. SEC fractions containing purified protein in DM were added to the lipid solution and incubated at RT for 25 minutes, then detergent was removed with Amberlite XAD-2 as previously described (47). As a negative control, POPC lipids were put through a simulated reconstitution process without protein. Amberlite was removed from each sample via gravity column and uniform liposomes were obtained by extrusion through a 0.2 µM membrane using an Avanti MiniExtruder. All SSME data were acquired using a Nanion SURFE2R N1 instrument. Liposome aliquots were thawed, diluted 2-fold, and briefly sonicated. 10 μL of liposomes were used to prepare 3 mm sensors as previously described(26). Before experiments, sensor capacitance and conductance values were obtained to ensure sensor quality. For all experiments, buffers contained 50 mM MES, 50 mM MOPS, 50 mM bicine, 100 mM NaCl, and 2 mM MgCl2 with internal pH values of 7.3 and external pH values of 7.0. For inward-facing drug gradients, external drug concentration was 8 μM and internal drug concentration was 0.5 μM. For outward-facing drug gradients, internal drug concentration was 8 μM and external drug concentration was 0.5 μM. Both internal and external drug concentration was 8 μM for the zero-gradient data. Sensors were rinsed with at least 500 μL of internal buffer before each measurement to set the internal buffer, pH, and drug concentrations as described in(26). Data acquisition occurred in three stages. First, sensors were perfused with an internal buffer, then transport was initiated by perfusion of the external buffer, and finally, perfusion of the internal buffer re-equilibrated the sensors. Signals were obtained by integrating the current during perfusion of the external buffer, with the final 100 ms of the initial internal buffer perfusion used as the baseline. Reported data are average values of at least three sensors, with error bars representing the standard error of the mean.

### Checkerboard Assays

MG1655 Δ*emrE E. coli* cells expressing either WT- or E14Q-EmrE constructs were grown in Mueller-Hinton broth (Sigma, 100µg/mL carbenicillin, pH 7.0) from a single colony to an OD of 0.2 at 37 °C. Kanamycin or tetracycline was serially diluted across a 96-well microplate in MHB with concentrations ranging from 0-80μM or 0-16μM respectively. Harmane was serially diluted down a separate plate using MHB with concentrations ranging from 0-1150μM. The cells were then diluted to a final OD of 0.01 in the microplate. A column with no kanamycin or tetracycline and a row with no harmane was used to determine the MIC values for each compound. Inoculated plates were sealed and incubated with shaking for 18h at 37°C. OD_600_ endpoints were taken using a microplate reader (BMG-Labtech). Checkerboard synergy testing was performed in triplicate and analyzed for MIC and FIC values in Excel.

The fractional inhibitory concentration (FIC) index was calculated for each well with no turbidity along the interface using the MIC values for the different compounds individually and in tandem. The MIC value was defined as the minimum concentration required to inhibit all cell growth to 10% of the background growth, as detailed in (29). FIC values were determined using the following equations:

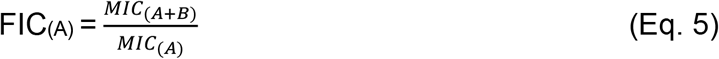

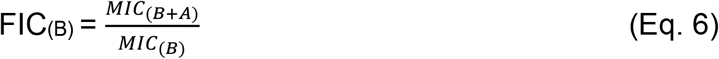

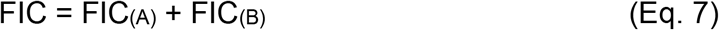

where A and B represent the different compounds in the assay. The mean FIC index was calculated and used to determine synergism (FIC < 0.5), indifference (0.5 ≤ FIC < 1), or antagonism (FIC ≥ 1).

### Bicarbonate Assays

MG1655 Δ*emrE E. coli* cells expressing either WT or E14Q-EmrE were grown overnight at 37 °C from a single colony. Harmane was serially diluted across a 96-well microplate in Mueller-Hinton broth (Sigma, 100µg/mL ampicillin, pH 7.0) from 0-0.4mM, with or without 25mM bicarbonate, and assayed with a starting OD_600_ of 0.01. Plates were then sealed and incubated with shaking for 18h at 37°C. OD_600_ endpoints were taken using a microplate reader (BMG-Labtech). Relative growth was calculated by dividing the measured OD_600_ from a given concentration by the OD_600_ for cells containing no drug. IC_50_ curves were performed in triplicate with three biological replicates per plate. Data was fit to a simple sigmoid equation using Igor Pro v8 (WaveMetrics Inc.).

## Acknowledgements

We thank Trey Sato for his assistance in using the OmniLog Instrument for Biolog Assays with funding from the Great Lakes Bioenergy Research Center under the U.S. Department of Energy, Office of Science, Office of Biological and Environmental Research Award Number DE-SC0018409. We also thank Andrea Wegrzynowicz for supplying the western blot expression data in Figure S1C. This study made use of the National Magnetic Resonance Facility at Madison, which is supported by NIH grants P41GM103399 (NIGMS) and R24GM141526 (NIGMS). This work was funded by NIH grants R01GM095839 and R35GM141748.

## Author contributions

Conceptualization – PJS, NET, KAHW; Methodology – NET, PJS, WFB, MB, CJP; Formal Analysis – PJS, NET, MB, KAHW; Validation – PJS, NET, MB, KAHW; Investigation – PJS, NET, WFB, MB, CJP, KMH; Writing (original draft) – PJS, NET, KAHW; Writing (review and editing) – PJS, NET, WFB, MB, CJP, KMH, KAHW; Visualization – PJS, NET, MB, KAHW; Supervision – PJS, KAHW; Funding acquisition – KAHW.

## Data Availability

Supplemental data and dataset information can be found in the SI of this paper.

