## Supplemental Information for "Activating alternative transport modes in a multidrug resistance efflux pump to confer chemical susceptibility"

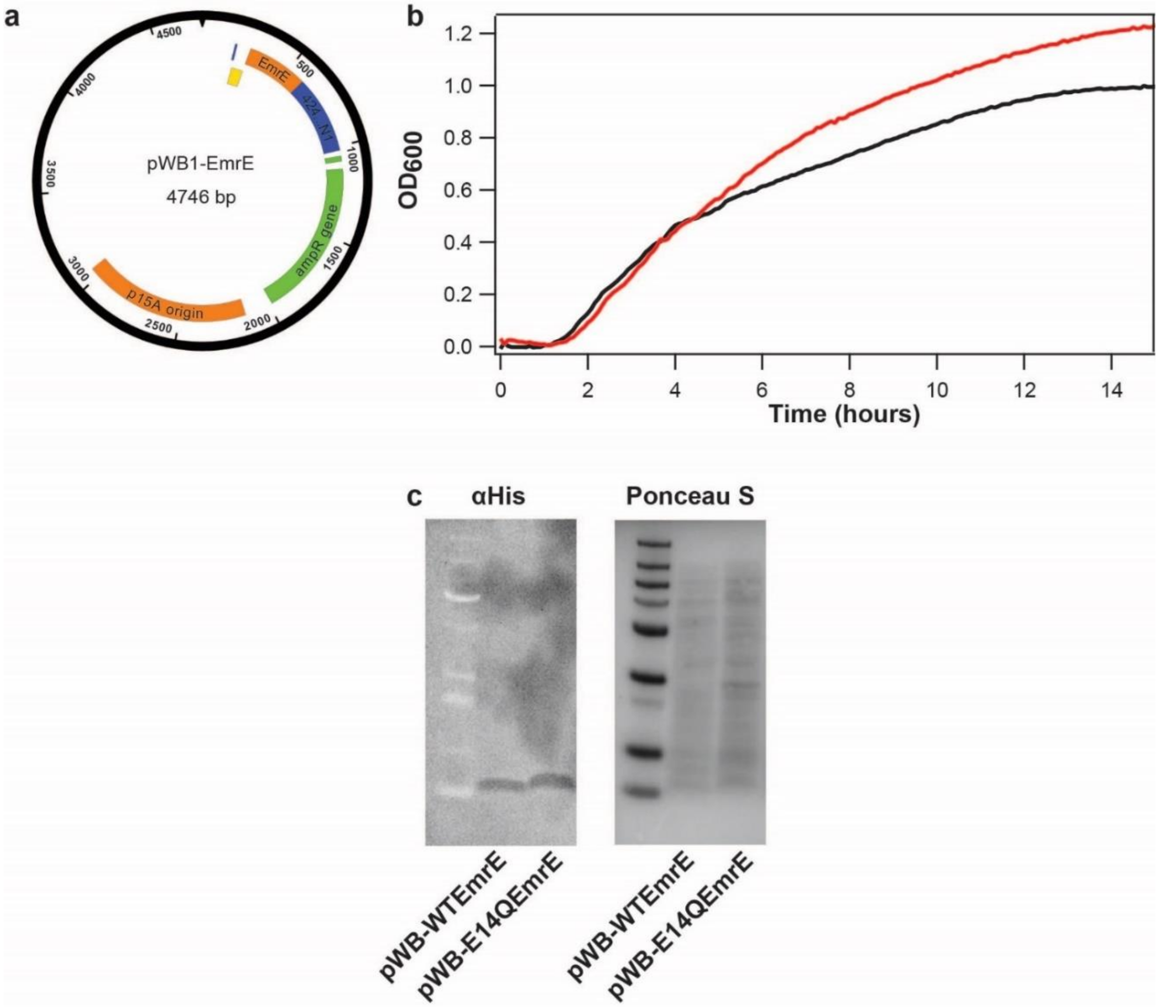


**Fig. S1. Leaky expression of WT-EmrE and non-functional, E14Q-EmrE have equal expression levels and similar growth patterns.** (a) The pWB plasmid expresses EmrE (WT or E14Q) under the control of a pTrc promoter. All *in vivo* experiments were carried out using only leaky expression of EmrE without induction by IPTG. (b) MG1655 Δ*emrE* cells expressing WT- or E14Q-EmrE were grown in Mueller-Hinton Broth with ampicillin in the absence of drug treatment. Note that E14Q-EmrE allows cells to grow to a higher $OD_{600}$ over time due to the inability to leak protons as WT-EmrE does. (c) Anti-His detection of leaky expression of WT- and E14Q-EmrE demonstrates that protein levels are equal for both constructs. Ponceau S stain of the western blot acts as a loading control.

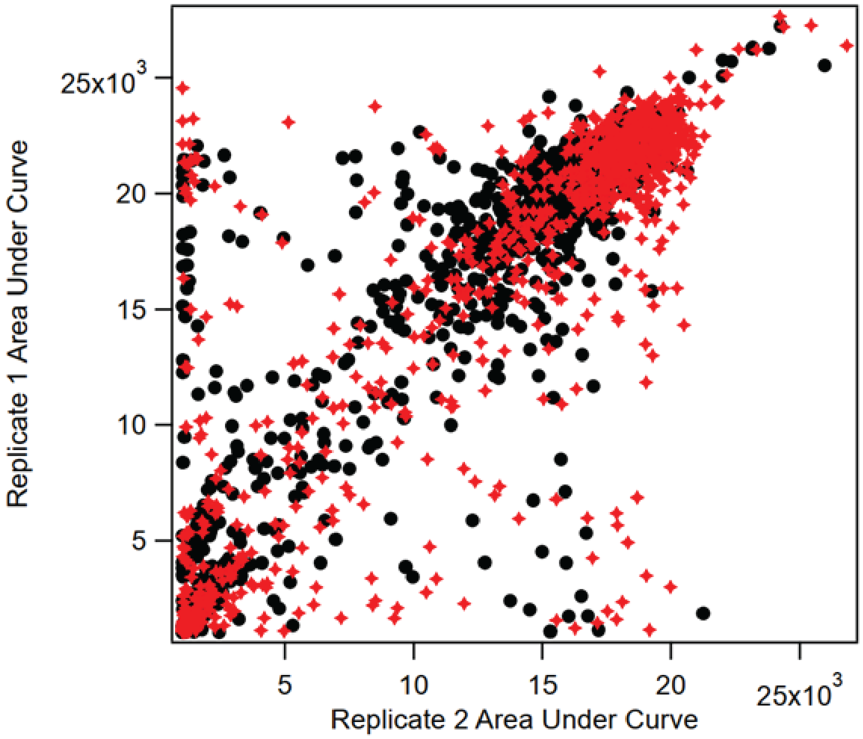

**Fig. S2. Two independent Biolog replicates are strongly correlated.** To quantify the correlation between replicates of the Biolog phenotype microarray, the area under the curve for all data in both biological replicates of the phenotype microarray for both WT-EmrE (black circles) and E14Q-EmrE (red diamonds) were plotted. The correlation constant for these data is 0.85 for WT and 0.84 for E14Q. These data were the basis for the hit threshold determination for results from EmrE data. Divergence in the data may stem from plate-to-plate differences in initial OD of cells, final compound concentration, and volume of sample.

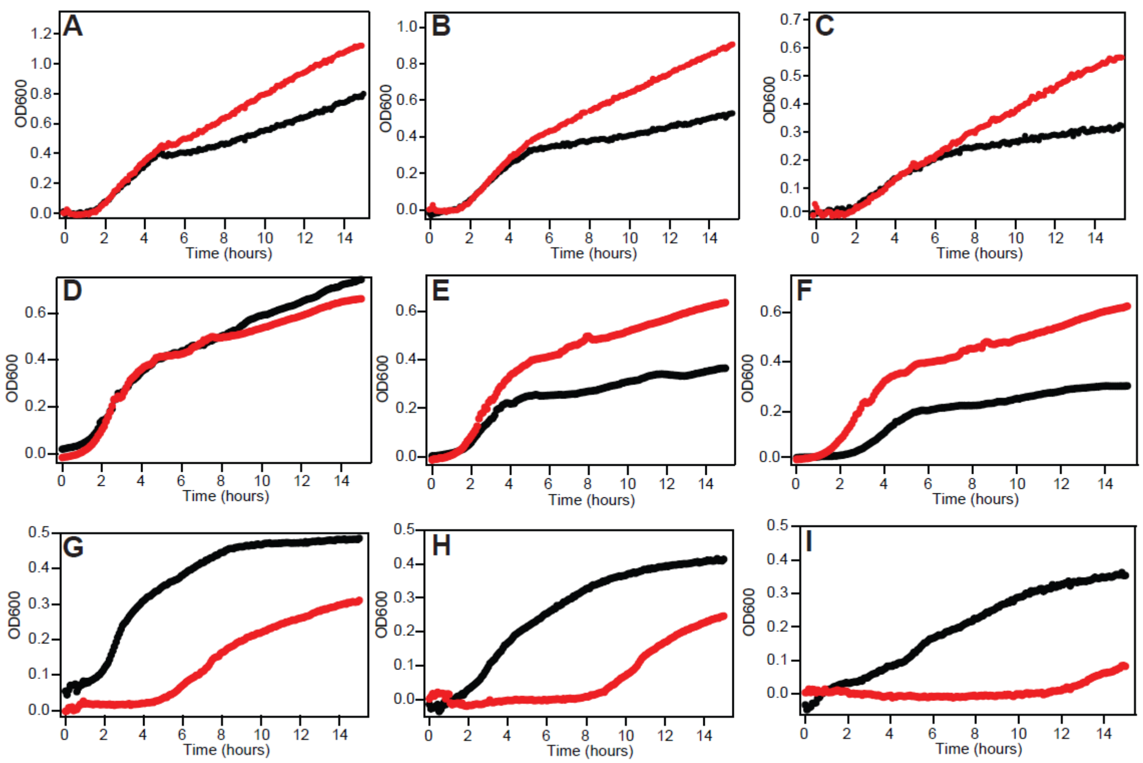


**Fig. S3. Growth curve phenotypes are dose-dependent.** To further validate the results of the Biolog phenotype microarray, growth curves at varying concentrations of harmane (A, 0.0325 mM; B, 0.065 mM; and C, 0.13 mM), 18-crown-6-ether (D, 0.1 mM; E, 0.6 mM; and F, 3 mM), and chelerythrine chloride (G, 0.2 mM; H, 0.4 mM; and I, 0.8 mM) were performed on MG1655 Δ*emrE* cells expressing WT-EmrE (black) or E14Q-EmrE (red). In all cases, the expected phenotypes (susceptibility for harmane and 18-crown-6-ether, and resistance for chelerythrine chloride) were observed with increasing compound concentration. Concentrations of compounds were selected based on the concentrations used in the Biolog phenotypic microarrays. The curves shown are an average of four biological replicates.

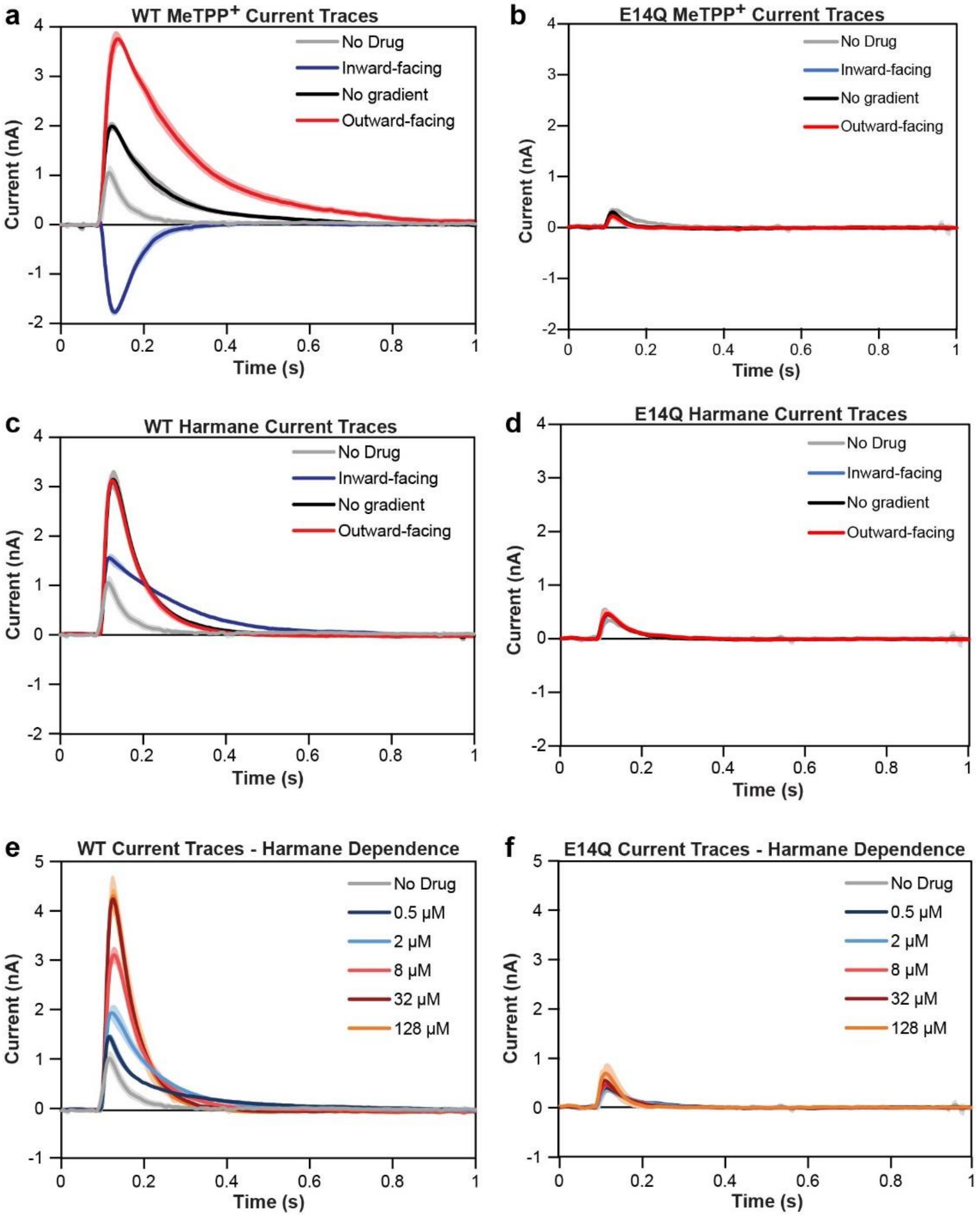


**Fig. S4. Raw current traces for SSME data.** Data shown is an average of at least three replicates, with standard deviation indicated by the shaded region. With both substrates, the minimal current is observed for sensors prepared using the non-functional mutant E14Q-EmrE (b, d, and f). $MeTPP^+$ (a and b) behaves as expected for an antiported substrate, with an increased signal when the drug and proton gradients are oriented in opposite directions and a reversal of transport direction when the large $MeTPP^+$ gradient is oriented in the same direction as the smaller proton gradient. Harmane (c and d) increases the transport signal compared to a background without drug. The signal is in the direction of downhill proton transport, regardless of the direction of the harmane gradient. Peak current of downhill proton transport increases in WT-EmrE proteoliposomes (e) with increased harmane concentration. E14Q-EmrE proteoliposomes do not show the same dependence on harmane concentration (f).

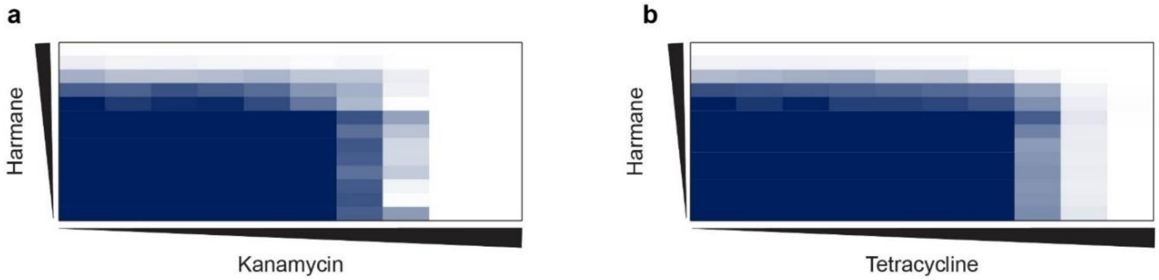


**Fig. S5. E14Q-EmrE checkerboard assays.** a) Checkerboard map of E14Q-EmrE expressing cells co-treated with harmane and kanamycin. b) Checkerboard map of E14Q-EmrE expressing cells co-treated with harmane and tetracycline. Concentration gradients are denoted by the black triangles. Growth is shown as a gradient of highest $OD_{600}$ (navy) to no growth (white).

**Table S1.** List of Biolog hits displaying resistance or susceptibility phenotypes.

| Compound | Hit Score | Formal Charge | Notes |
|---|---|---|---|
| Harmane | -6 | 0 | Specific reversible inhibitor of monoamine oxidase A |
| Hexachlorophene | -6 | 0 | Antifungal, antiseptic |
| Menadione | -5 | 0 | Vitamin K |
| 18-crown-6-ether | -5 | 0 | Metal-binding |
| Cefoperazone | -4 | 0 | Cephalosporin antibiotic |
| Nitrofurazone | -4 | +1 | Antibiotic, interference of DNA synthesis |
| Oxytetracycline | -4 | 0 | Antibiotic |
| Cobalt (II) chloride | -4 | 0 | metal |
| Spectinomycin | -4 | 0 | Antibiotic |
| Ethionamide | -4 | 0 | Prodrug antibiotic, |
| Rolitetracycline | -3 | 0 | Antibiotic, protein synthesis inhibition |
| Geneticin disulfate | -3 | 0 | Antibiotic, protein synthesis inhibition |
| Ruthenium red | -3 | 0 | Inorganic dye |
| Antimony (III) chloride | -3 | 0 | Metal for vitamin A detection |
| Troleandomycin | -3 | 0 | Macrolide antibiotic |
| Cefoxitin | -3 | 0 | Cefalosporin antibiotic |
| Coumarin | -3 | 0 | Metabolite |
| Nickel chloride | -3 | 0 | Metal |
| Oleandomycin | -3 | 0 | Macrolide antibiotic |
| Erythromycin | -3 | 0 | Macrolide antibiotic |
| Dodine | -3 | 0 | Fungicide |
| Glycine HCl | -3 | 0 | Non-essential amino acid |
| Spiramycin | -3 | 0 | Macrolide antibiotic |
| Manganese (II) chloride | 3 | 0 | Metal |
| Methyltrioctylammonium chloride | 3 | +1 | Phase transfer catalyst |
| FCCP | 3 | 0 | Proton ionophore |
| Tetrazolium violet | 3 | +1 | Dye, apoptosis inducer, antineoplastic agent |
| Cetylpyridinium chloride | 4 | +1 | Antiseptic, destabilizes cell membranes |
| Acriflavine | 4 | +1 | Local antiseptic, biological stain |
| Sanguinarine chloride | 4 | +1 | Toxic polycyclic ammonium ion |
| Proflavine | 4 | 0 | Bacteriostat |
| Chelerythrine chloride | 5 | +1 | Antibiotic, apoptosis induction |
| Crystal violet | 6 | +1 | Topical antibiotic |
| Methyl viologen | 8 | +2 | Herbicide, desiccant, photosystem-I inhibitor |

**Table S2.** Buffer Conditions for SSME experiments.

| **Drug Gradient** | **Internal Buffer** | **External Buffer** |
|---|---|---|
| Inward-facing | 0.5 µM drug<br>50 nM $H^+$ (pH 7.3) | 8 µM drug<br>100 nM $H^+$ (pH 7.0) |
| Outward-facing | 8 µM drug<br>50 nM $H^+$ (pH 7.3) | 0.5 µM drug<br>100 nM $H^+$ (pH 7.0) |
| No gradient | 8 µM drug<br>50 nM $H^+$ (pH 7.3) | 8 µM drug<br>100 nM $H^+$ (pH 7.0) |
| No drug | 50 nM $H^+$ (pH 7.3) | 100 nM $H^+$ (pH 7.0) |
